# *Candida albicans* cells lacking AP-2 have defective hyphae and are avirulent despite increased host uptake and intracellular proliferation in macrophages

**DOI:** 10.1101/2023.08.03.551781

**Authors:** Stella Christou, Kathryn R. Ayscough, Simon A. Johnston

**Author notes:** corresponding author Simon A Johnston <. KRA and SAJ are joint senior authors.

## Abstract

*Candida albicans* is a commensal microbe and opportunistic human pathogen. The yeast can be recognised and taken up by macrophages via interactions with its cell wall, a complex polysaccharide structure containing several components that are specifically recognised by immune cell receptors. Following uptake *Candida* can respond in the host environment by switching from a yeast to hyphal morphology which facilitates escape from macrophages and allows subsequent invasion of host tissues. Disruption of *Candida*’s ability to form hyphae results in reduced virulence and fitness for survival in the host environment. *Candida albicans* cells lacking AP-2, an endocytic adaptor complex, have increased cell wall chitin and morphologically defective hyphae *in vitro*. Previous studies have correlated increased chitin with decreased recognition by macrophages, possibly due to masking of cell wall beta-glucan which is recognised by dectin receptors. Despite the cell wall changes in the mutant strain there was an unexpected increased uptake of the mutant. Increased chitin did not reduce phagocytosis and additional uptake was not due to compensatory elevated exposure of beta-glucan, highlighting the importance of cell wall components beyond chitin and glucan for macrophage engagement and uptake. Furthermore, the *apm4* mutant exhibited parasitism of macrophages, surviving and proliferating within the phagosome, a phenotype that was then replicated with a well-characterised yeast locked mutant.

Finally, the combined phenotype of reduced hyphal formation but continued proliferation resulted in reduced virulence despite an equivalent burden of infection to a wild-type *Candida* infection, as determined using a zebrafish larval model of candidiasis.

## Introduction

Candida albicans is a commensal microbe and opportunistic pathogen of humans. Cellular innate immunity is essential for control of Candida infection and *C. albicans* cells are recognized and taken up by macrophages (Hernandez-Chavez et al., 2017; Newman and Holly, 2001; Scherer et al., 2020). The *C. albicans* cell wall is a complex polysaccharide structure that contains several chemical structures that are specifically recognized by the immune cell receptors e.g. β glucan binding and recognition by Dectin-1 (Gow et al., 2007; Hasim et al., 2017; Herre et al., 2004; Mansour et al., 2013). Chitin is another component of *Candida* cells walls and that has been implicated with virulence of Candida. Chitin levels and exposure have been correlated to both pro- and anti-inflammatory responses in immune cells. Elevated cell wall chitin in *Candida* leads to increased resistance to the anti-fungal drug caspofungin and chitin appears to disrupt recognition of β -glucan uptake by macrophages (Mora-Montes et al., 2011). In addition to the molecular disruption of the immune response to infection, Candida can respond to the host environment by switching from a yeast to hyphal morphology, a strategy used to escape the macrophage phagosome (Vylkova et al., 2011; Vylkova and Lorenz, 2014; Westman et al., 2018). Not only can the formation of hyphae support escape from macrophages but hyphal cells can also aid the invasion of host tissues (Moyes et al., 2016; Newman and Holly, 2001; Seman et al., 2018; Swidergall, 2019). Disruption of *C. albicans* cells’ ability to form hyphae results in reduced virulence and fitness for survival in the host environment (Lo et al., 1997; Scherer et al., 2020; Seman et al., 2018).

We have previously demonstrated that deletion of the gene encoding the mu subunit (*apm4*) of the heterotetrametric AP-2 endocytic adaptor complex in *C. albicans* leads to an inability to form a stable and functional complex at the plasma membrane. A major endocytic cargo that has been identified for *C. albicans* AP-2 is Chs3, the major chitin synthase in *C. albicans*. The *apm4* deletion does not affect secretion of Chs3 but does lead to inhibition of its endocytosis resulting in elevated levels of Chs3 at the cell surface and increased levels of chitin in the cell wall (Knafler et al., 2019). Despite the cell wall changes, the *apm4Δ/Δ* strain can undergo a hyphal switch in appropriate conditions but the hyphae that form are shorter and wider. Heterozygous deletion of *chs3* in the *apm4Δ/Δ* strain, results in a reduction of chitin on the cell surface as expected, and also leads to a partial rescue of the hyphal morphology phenotype demonstrating the importance of chitin levels and its distribution for facilitating appropriate cell shape (Knafler et al., 2019). Given the phenotype of the *apm4* mutant we hypothesized that its virulence in macrophages and during invasive infection would be reduced due to its defect in formation of hyphae, but that increased cell wall chitin might enhance evasion of macrophages. Therefore, we sought to determine how the combined phenotypes of *apm4Δ/Δ* cells influenced the interaction of Candida with macrophages and the outcome of infection. Studying the interaction with macrophages, we revealed an unexpected increased uptake of the *apm4Δ/Δ* mutant. Dissection of this phenotype revealed increased chitin did not reduce phagocytosis and this was not due to compensatory elevated exposure of beta-glucan, highlighting the importance of cell wall components beyond chitin and glucan for macrophage engagement and uptake. Furthermore, despite its inability to form hyphae inside macrophages the *apm4* mutant still exhibited parasitism of macrophages, both surviving and proliferating within the phagosome, a phenotype that we then replicated with a well characterised yeast locked mutant, NRG1. Finally, using a zebrafish larval model of candidiasis we determined that the combined phenotype of reduced hyphal formation but growth within macrophages resulted in reduced virulence despite an equivalent burden of infection to wild type Candida infection.

## Results

### Apm4 deletion results in increased uptake by macrophages despite increased levels but similar beta-glucan exposure

Deletion of *apm4* has been shown to result in an increase in the Candida cell wall chitin levels (Knafler et al., 2019). To determine whether the cell wall changes in the *apm4Δ/Δ* mutant influence uptake by murine macrophages, adhered murine J774 macrophage-like cells were infected with wild type and *apm4Δ/Δ C. albicans*. We found that the *apm4Δ/Δ C. albicans* cells were phagocytosed by a greater proportion of macrophages (1.3-fold increase; Figure 1A,B) and that each macrophage on average phagocytosed a larger number of yeast cells (Figure 1C). This finding indicated that either the increased chitin exposure did not reduce phagocytosis of *apm4Δ/Δ C. albicans* or that there was an additional phenotype that was counter-acting the effect of increased chitin in the cell wall on phagocytosis. We hypothesized two possible counter-acting factors: that mutant cells were smaller which would facilitate more rapid phagocytosis or that β-glucan levels were increased in *apm4Δ/Δ C. albicans*. First, we measured length and width of wild type and mutant cells and, in contrast to our hypothesis, found that the *apm4Δ/Δ C. albicans* were larger than wide type cells (Figure 1D). *C. albicans* are typically ellipsoid in their yeast form and this was unaltered in the *apm4Δ/Δ* mutant. Calculation of surface area based on length and width data indicated a surface area of 69 μm2 for wild type cells compared to 111 μm^2^ for the mutant. Therefore, the *apm4*Δ/Δ mutant presents a significantly greater surface area available for interaction with macrophages.

**Figure 1:**
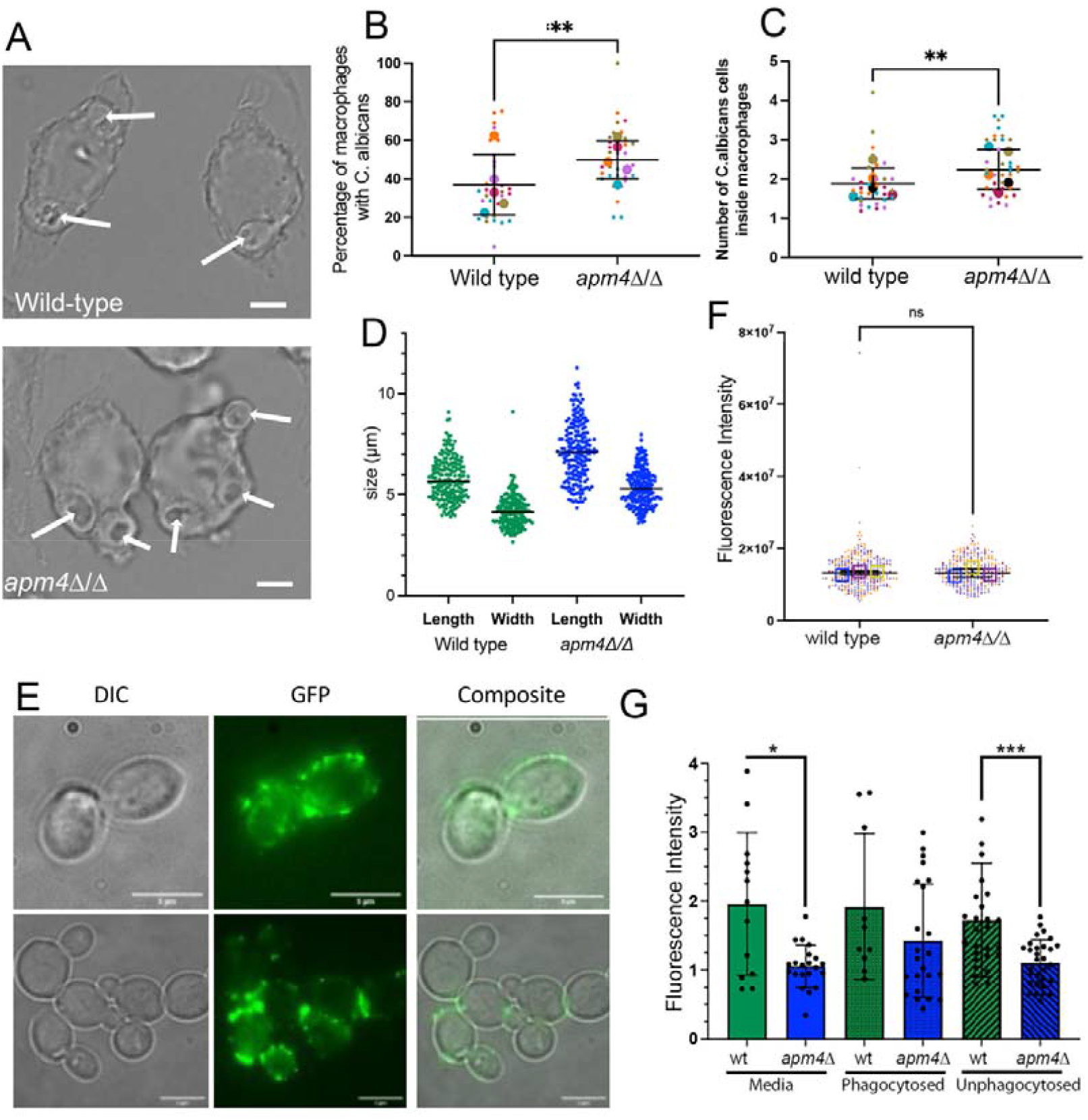
Deletion of *apm4* in *Candida albicans* results in increased levels of phagocytosis but exposed β-glucan is unchanged. *C. albicans* and J774 macrophages were co-incubated at 1:10 ratio for 30 minutes at 37°C, then fixed and imaged. (A) Wild type and *apm4*Δ*/*Δ infected macrophages. White arrows indicate phagocytosed C. albicans; scale bar 5 μm. B) The proportion of macrophages in a field of view that phagocytose C. albicans were quantified. 5 independent replicates with approximately 6 images per replicate. Larger circles represent the average of each replicate, and small circles represent the values for each image analyzed. Mann-Whitney statistical tests show significance, p value 0.008. C) The number of *C. albicans* per macrophage. 5 independent replicates with 6 images per replicate. Larger circles represent the average per replicate, small circles represent the values for each image analyzed. Mann-Whitney statistical tests show significance with p value 0.0072. *D)* Overnight cultures of C. albicans were refreshed with YPD media for 30 minutes and then images of cells were captured and analyzed for cell size. n=200 for each strain., p<0.0001 for both length and width comparison between wild type and mutant. The levels of exposed β-glucan were investigated using immunofluorescence microscopy. (E) Overnight cultures of *C. albicans* were incubated with fresh media for 3 hours before fixing and staining with Fc-Dectin. (F) Fluorescence intensity of the cell staining was quantified as described. Shown is a SuperPlot of exposed β glucan from 3 biological replicates. (G) J774 macrophage-like cells were infected with Candida albicans for 30 minutes before fixation, then stained and imaged for exposed Fc-Dectin. The images captured were used to analyse levels of Fc-Dectin in phagocytosed and non-phagocytosed *C. albicans* and compare that to *C. albicans* cells that are incubated in media without J774 cells. The data represent the fluorescence intensity of the cells. * p =0.0321, *** p = 0.0004

Both total β-glucan levels and its exposure at the cell surface are recognized as important factors affecting uptake of *C. albicans* cells by macrophages (Bain et al., 2014; Munro, 2013). The total levels of β glucan in *apm4Δ/Δ* cells were previously reported to show no overall change compared to wild type cells (Knafler et al., 2019). However, given the increased phagocytosis of mutant cells we investigated if there was greater surface exposure of β-glucan and this could be the factor responsible for the increased uptake and recognition by the macrophages (Davis et al., 2014; Galan-Diez et al., 2010; Gow et al., 2007; Hasim et al., 2017; Mora-Montes et al., 2011). For wild type cells the expected pattern of exposed β glucan in patches around the cell surface and near the bud neck regions was observed and this distribution was replicated in the absence of *apm4* (Figure 1E). Furthermore, there was no significant difference in the exposed β-glucan levels between the two populations when fluorescence intensity of staining was quantified (Figure 1F). The results indicate that the changes in the cell wall composition of *apm4Δ/Δ* cells do not result in a change of the exposed β-glucan levels and suggested that the increased phagocytosis is not linked to β-glucan recognition. However, we also considered the possibility that β-glucan levels could become altered when cells were exposed to mammalian cell media or in the presence of macrophages and that this resulted in increased exposure of β-glucan on *apm4Δ/Δ* versus wild type cells. To address this, the levels of exposed β-glucan were measured during interactions with macrophages. However, in contrast to our hypothesis the levels of exposed β-glucan were decreased in extracellular *apm4Δ/Δ* cells in both media alone and in the presence of macrophages.

There was no significant difference however, for either wild type cells or *apm4Δ/Δ* when compared across conditions (Figure 1G). The data indicate that as expected, the increased chitin in the *apm4Δ/Δ* mutant correlates with decreased exposure of β glucan. However, the increased chitin in *apm4Δ/Δ C. albicans* did not lead to reduced uptake by macrophages and that factors other than β-glucan and cell size are responsible for increased phagocytosis.

### Intracellular *apm4Δ/Δ C. albicans* have a defect in hyphal formation in macrophages

Our previous *in vitro* study of the *apm4Δ/Δ C. albicans* demonstrated a defect in the formation of hyphae. Heterozygous deletion of the major chitin synthase *chs3* in the *apm4Δ/Δ* background however, reduced cell wall chitin levels and partially rescued the ability of mutant cells to form true hyphae demonstrating an important and specific role for chitin in the hyphal deficiency phenotype (Knafler et al., 2019). Following phagocytosis, wild-type *C. albicans* cells switch morphology, forming hyphae that can rupture the plasma membrane so lysing the host macrophage (El-Kirat-Chatel and Dufrene, 2012; Hernandez-Chavez et al., 2017; Westman et al., 2018; Westman et al., 2020). We hypothesized that despite increased uptake into macrophages, intracellular *apm4Δ/Δ C. albicans*, would be unlikely to lyse the cells due to the defect in hyphal switching. Using time lapse microscopy, we measured the formation of hyphae by intracellular Candida and the lysis of macrophages. While hyphal Candida were seen in nearly all fields of view following infection of macrophages with wild type cells, they were rare following phagocytosis of *apm4Δ/Δ C. albicans* (Figure 2A). To quantify this difference, we counted the number of yeast-form (including budding cells) and hyphal-form (both true and pseudo-hyphae) cell morphologies over 18 hours of infection (Figure 2B). As expected, the majority of intracellular wild type Candida cells (>80%) switched from a yeast to hyphal morphology over the 18 hours of imaging (Figure 2C). In contrast, most *apm4Δ/Δ* cells remained in the yeast form. Hyphal-like cells were observed in a proportion of macrophages infected with the *apm4* mutant, but no elongated hyphae or penetration of macrophages were observed. We hypothesized that the *apm4Δ/Δ, chs3Δ/CHS3* strain would be better able to form hyphae in macrophages, as had previously been demonstrated *in vitro*. In agreement with our hypothesis, we found that reduced chitin levels in this strain resulted in an increase in the proportion of yeast cells that underwent hyphal switching. Thus, despite increased phagocytosis by macrophages and increased levels of cell wall chitin, the *apm4Δ/Δ* cells were less able to evade macrophages by intracellular hyphal switching.

**Figure 2:**
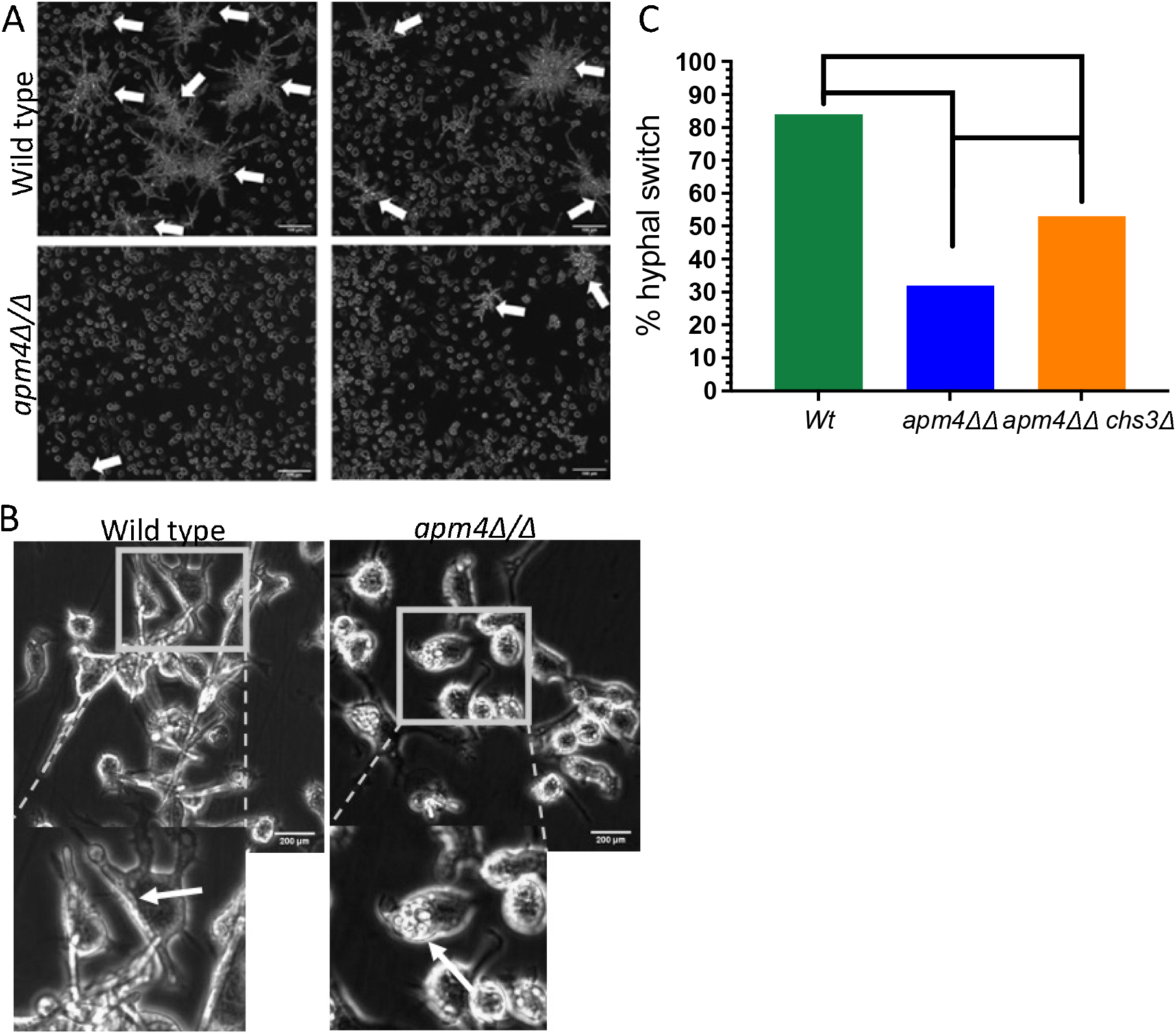
*Candida albicans* cells with an AP-2 deletion show hyphal switch deficiencies. *C. albicans* and J774 macrophages at 1:1 ratio were co-incubated for 18 hrs. The interacting cells were imaged at 20x and they were analysed for hyphal morphology switching. (A) Representative images of infection after 18 hours of interaction are shown. The white arrows indicate macrophage, C. albicans co-clusters. Scale bars are 100μm. (B) Time-lapse captures at 10 hours post interaction. The panels focus on individual macrophage cells, the white arrow illustrate phagocytosed C. albicans. (C) The timelapses were analyzed by single cell observations for hyphal switch in the phagosome from 75 infected macrophages. The graph represents the quantification of hyphal switch inside the phagolysosome. Cell morphology changes that occurred only inside the phagolysosome were recorded.

### Intracellular proliferation of yeast form Candida *cells* in the absence of hypha formation

Given the inability of *apm4Δ/Δ* mutant cells to lyse macrophages by the formation hyphae we predicted that macrophages would digest intracellular *apm4Δ/Δ* Candida. Using time lapse microscopy, we quantified the fate of intracellular *apm4Δ/Δ* mutant cells. In contrast to our prediction, we found that the mutant cells were not digested but instead were able to proliferate inside macrophages (Figure 3A,B). Remarkably, over 40% of the *apm4Δ/Δ* cells that were phagocytosed were able to proliferate inside the phagosome while fewer than 10% of the wild-type population showed this behaviour. To determine whether proliferation was a switching response to the failure to maintain hyphal growth we tested the a*pm4Δ/Δ, chs3Δ/+* cells. We predicted that the increase in hypha formation would proportionally reduce the cells that proliferated inside macrophages. In agreement with our prediction, we measured an almost identical reduction in budding compared to the increased rate of hyphal formation in a*pm4Δ/Δ, chs3Δ/+* cells (Figure 3B).

**Figure 3:**
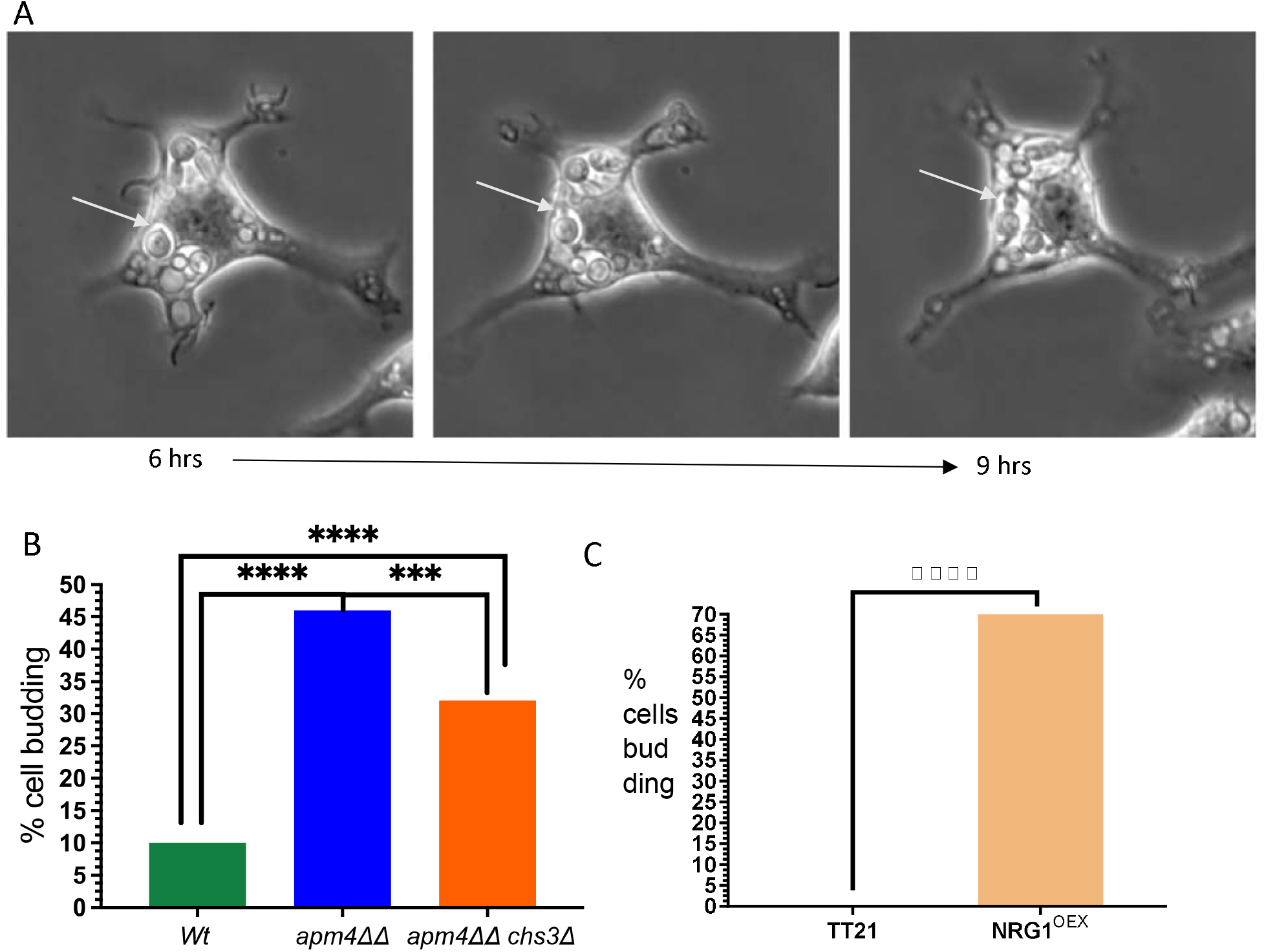
*Candida albicans* cells with *apm4* deletion grow inside the phagolysosome. *C. albicans* and J774 macrophages at 1:1 ratio were co-incubated for 18 hours. The interaction was recorded via timelapse microscopy. Timelapses were then analyzed by single cell observations. A) Timelapse images from the interaction of J774 cells with *C. albicans* over time. The white arrows show *C. albicans apm4Δ/Δ* cells budding inside the host. The data were analysed for the percentage of cells forming buds inside the phagosome in(B) macrophages are infected with Wt, *apm4Δ/Δ and apm4Δ/Δ, chs3Δ/CHS3* and (C) macrophages infected with TT21 and the yeast locked strain NRG1^OEX^. Fisher’s exact test was used to analyse the statistical significance of the results.

We next sought to establish whether the capacity to proliferate vegetatively in the phagosome was conserved in other mutants with a defect in formation of hypha. We chose the NRG1^OEX^ mutant that does not form hyphae but through a different mechanism to *apm4Δ/Δ* cells. While *apm4Δ/Δ* mutant cells are able to initiate the formation of hyphae their defect in endocytic recycling at the hyphal tip disrupts the progression of hyphal growth (Knafler et al., 2019), in contrast the NRG1^OEX^ mutant is unable to initiate hyphal formation (Murad et al., 2001). The parental strain (TT21) formed hyphae in the macrophages and no budding cells were observed (Figure 3). In contrast, the majority of NRG1^OEX^ cells proliferated inside macrophages and at a greater proportion than the *apm4Δ/Δ* cells, as might be expected by the more severe hyphal switch defect in NRG1^OEX^ cells. Therefore, we could demonstrate that two different Candida mutants with defects in hyphal formation were able to use alternative macrophage parasitism strategies and proliferate.

### Impact of the *apm4* deletion on virulence in zebrafish

The data thus far revealed an intriguing phenotype of *apm4Δ/Δ* cells in their interaction with macrophages. While unable to escape macrophages by formation of hyphae, *apm4Δ/Δ* cells were able to still avoid killing and were observed to proliferate within macrophages. The decrease in formation of hyphae in *C. albicans* is well evidenced to reduce its virulence in animal models of *Candida* infection (Lo et al., 1997; Seman et al., 2018). However, the balance of macrophage parasitism by intracellular proliferation versus cell and tissue damage by hyphae is less well understood. We used a zebrafish larval model of Candida infection because we were able to directly visualise the Candida phenotype in vivo while simultaneously measuring the progression of infection. Blood-stream infection of zebrafish embryos 1 day post fertilisation with wild type *C. albicans* cells resulted in the presence of numerous hyphal candida 1 day post infection (dpi) (Figure 4A). In contrast, as we had previously shown in vitro and following phagocytosis by macrophages, *apm4Δ/Δ* mutant cells were deficient in the formation of hyphae. Comparison of fungal burden between wild type and mutant cells was challenging because of the large difference in the populations of hyphal and yeast candida between the two strains. We hypothesised that the *apm4Δ/Δ* mutant cells would proliferate in zebrafish in their yeast form as we had shown inside macrophages. We decided to use measurement of fungal burden in vivo in zebrafish using the fluorescence intensity of GFP-tagged strains because this allowed the comparison of combined hyphal and yeast burden. Using this method we predicted that there would be no difference in fungal burden as the increased fluorescence of hyphal Candida would be matched by the increased number of yeast-form *apm4Δ/Δ* mutant cells. Measurement of fungal burden 1 and 2 dpi demonstrated an increase over time but no difference in total fluorescence between wild type and *apm4Δ/Δ* infections (Figure 4B). Having demonstrated a similar phenotype of *apm4Δ/Δ* mutant cells *in vivo* in zebrafish in comparison to infection of macrophages, we sought to identify any differences in virulence between hyphal dominated wild type infections and proliferating yeast dominated *apm4Δ/Δ* infections. Infection of zebrafish embryos 1 day post fertilisation resulted in 70% mortality over 4 days of infection. In contrast, very few *apm4Δ/Δ* infected zebrafish succumbed to infection and we found less than 20% mortality 4 days post infection. Finally, using the intermediate phenotype *apm4 Δ/Δ chs3 +/Δ* mutant we demonstrated increased virulence compared to *apm4 Δ/Δ C. albicans* further demonstrating the importance of chitin and hyphal formation for the virulence phenotype.

**Figure 4:**
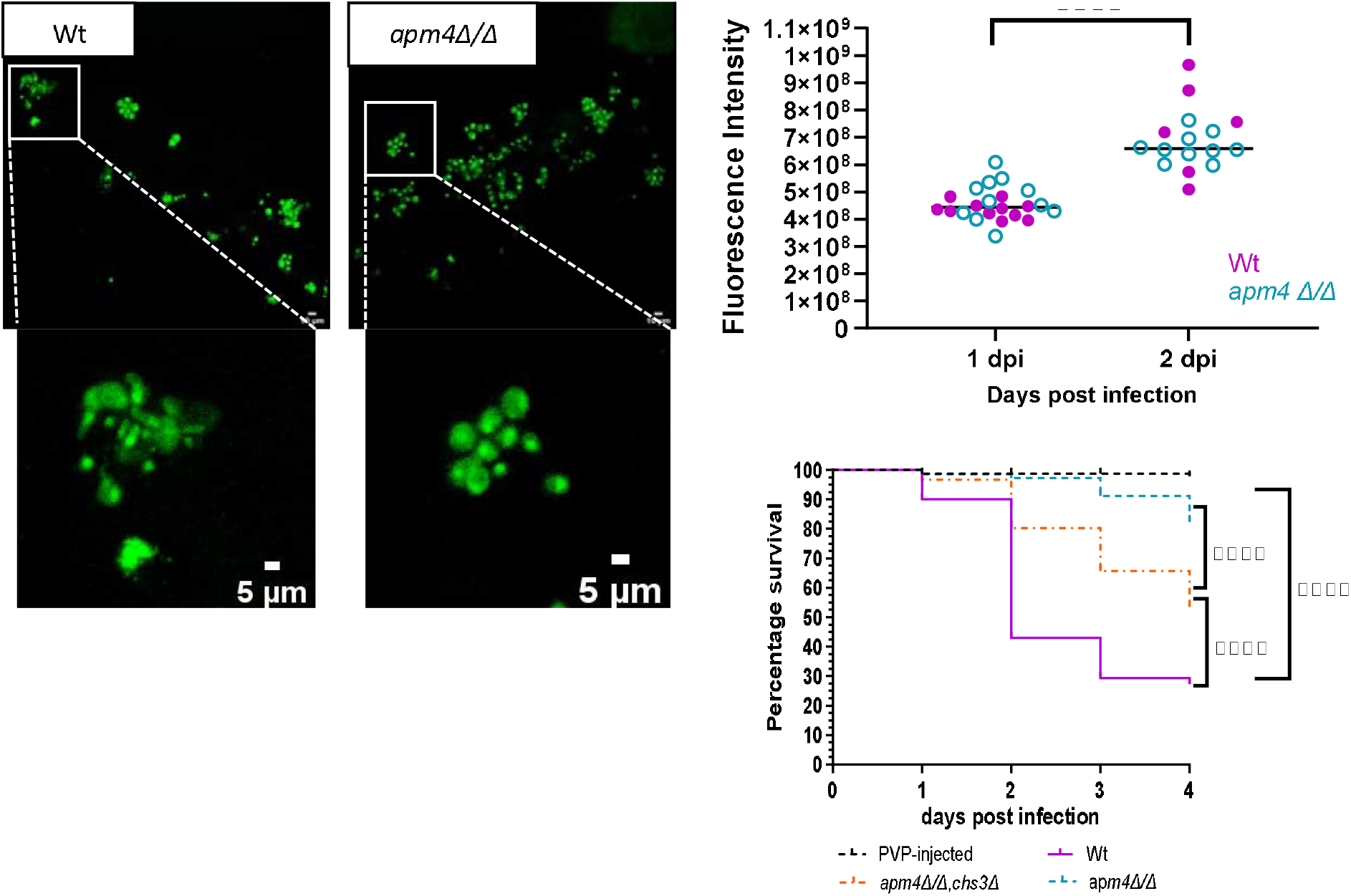
Candida albicans cells with apm4 deletion have a hyphal switch deficiency and reduced virulence during zebrafish embryo infections. Nacre zebrafish embryos at 1 dpf were injected with 500 cfu of C. albicans in the caudal vein to monitor the infection progression. A) Confocal images of the infected embryos at 1 dpi were captured to observe the morphology of C. albicans cells in the zebrafish bloodstream. B) The fungal burden of embryos infected was assessed using fluorescence from 3 biological replicates. Embryos were imaged at 1 and 2 dpi. Candida cells injected into embryos expressed cytoplasmic GFP and thus th e fluorescence emitted during microscopy represented Candida cells. A Z-stack was captured for each embryo and the fluorescence intensity analysis was conducted using the Maximum Projection of the images. C) Embryos from 3 biological replicates were monitored for survival up to 5 dpf. Survival was assessed by cessation of heartbeat. The number of surviving embryos was counted every day until 4 dpi. The zebrafish were infected with the Wt, apm4Δ/Δ, apm4Δ/Δ, chs3Δ and the PVP as an injection control. Survival statistical analysis concluded p<0.0001 from an n=90 embryos/group.

## Discussion

### Phagocytosis is increased for *C. albicans* with an *apm4* deletion

The cell wall of *Candida albicans* has long been recognised as a key component in recognition and phagocytosis of the pathogen by macrophages. A significant body of research has demonstrated the importance of exposed β-glucan in the cell wall binding to dectin-1 receptors on macrophages as part of this uptake process (Davis et al., 2014; Galan-Diez et al., 2010; Hasim et al., 2017; Mansour et al., 2013). Conversely, increased chitin has been considered a way that Candida could mask β-glucan and thus avoid immune recognition (Mora-Montes et al., 2011; Munro, 2013). In our previous work we had shown that inhibition of the AP-2 endocytic adaptor function through deletion of one subunit of the complex (*apm4*), led to an increased level of the major chitin synthase Chs3 at the Candida cell surface. There was a corresponding increase in cell wall chitin (Knafler et al., 2019). It was therefore unexpected in this study when the *apm4Δ/Δ* cells showed increased uptake by macrophages especially as the β-glucan remained similarly exposed or slightly masked compared to wild-type cells. The results suggest that cell wall components other than β-glucan are more important for the increased uptake of *apm4Δ/Δ* cells. Chitin currently has no known host receptors involved in its recognition, but it is co-recognized with mannans (Netea et al., 2008; Snarr et al., 2017). Previous analysis of the cell wall of *apm4Δ/Δ* cells revealed an increase in the thickness of the mannan layer albeit with no overall proportional increase in mannan (Knafler et al., 2019). Together the data support the idea that mannans are a critical component in recognition by macrophages and that recognition is not reduced in the *apm4Δ/Δ* mutant despite increased chitin levels.

### The virulence of *C. albicans* cells with an *apm4* deletion is reduced

Following uptake by macrophages, a *C. albicans* cell protects itself from host cell defence mechanisms and undergoes a morphological change to form a long projection or hypha to bring about host cell lysis (El-Kirat-Chatel and Dufrene, 2012; Hernandez-Chavez et al., 2017; Westman et al., 2018; Westman et al., 2020). The hyphal morphology is also critical for tissue invasion and damage through the production of the *C. albicans* toxin Candidalysin (Moyes et al., 2016; Richardson et al., 2018). While in the macrophage, the pathogen induces host arginase to reduce production of nitric oxide a key component in the host cell defence (Wagener et al., 2017). The hyphal switch of *apm4Δ/Δ* cells was investigated during interactions with the macrophage phagosome. It was observed that *apm4Δ/Δ* cells have a hyphal switch deficiency, though many cells (∼30%) do initiate short hyphae. Despite being observed within the phagolysosome, the mutant cells however did not appear to result in marked macrophage killing. Reduction of chitin in the cell wall by deletion of one copy of *chs3* in the *apm4* mutant led to a rescue in the proportion of cells that could form hyphae within the phagolysosome indicating the importance of chitin levels for the phenotypes observed.

Virulence was also assessed during in vivo infections of zebrafish embryos where it was concluded that *C. albicans* cells with an *apm4* deletion are significantly less virulent than wild-type cells. *Candida albicans* strains that do not show a hyphal switch have previously been reported to be avirulent, with the yeast locked strains demonstrating this phenotype most clearly (Lo et al., 1997; Seman et al., 2018). A reduction in cell wall chitin in the mutant partially restored virulence re-enforcing evidence for the importance of chitin in formation of true hyphae and virulence (Lo et al., 1997; Seman et al., 2018).

### Survival and proliferation in the host

As well as undergoing hyphal switch, another property of Candida cells following uptake by macrophages is that they induce host arginase and this acts to protect the pathogen from host defence killing mechanisms involving production of nitric oxide (Wagener et al., 2017). This mechanism promotes Candida cell survival in the macrophage environment and has been considered a precursor to hyphal formation (Ghosh et al., 2009). In the case of the *apm4Δ/Δ* cells however, which cannot produce lysis-competent hyphae we could show that cells continue to survive within the phagosome as budding of cells could be observed. This might indicate that the cells are able to induce host arginase irrespective of an ability to grow or maintain hyphae. Wagener and colleagues also demonstrated that increased surface chitin can lead to an increase in host arginase and that this can lead to pathogen protection (Wagener et al., 2017). Elevated chitin in the *apm4* mutant could therefore be postulated to be part of the mechanism facilitating mutant survival in the phagosome. This however cannot be the entire explanation for apm4Δ/Δ survival as the continued budding of the yeast-locked NRG1^oex^ cells in the same environment would suggest that increased chitin is not pre-requisite for continued proliferation. Our results suggest that *Candida*, irrespective of their capacity to produce hyphae, are able to induce conditions in the phagosome which facilitate its own survival and protect it from host killing mechanisms

Continued survival and proliferation of *apm4Δ/Δ* Candida cells was also found in zebrafish. While wild-type Candida led to a high level of host killing, the *apm4Δ/Δ* mutant was able to continue to proliferate as judged by an increased fungal burden, but it did not lead to high levels of death. A similar finding has also been reported for continued proliferation of yeast-locked strains in zebrafish (Scherer et al., 2020).

As in the case of macrophages, restoration of cell wall chitin to wild-type levels in the *chs3*Δ heterozygote (Figure 4), correlated with the capacity to form normal hyphae and partially restored virulence.

Overall our data indicate the importance of an optimal level of cell wall chitin for the growth and maintenance of hyphae that are effective for macrophage lysis or tissue invasion in vivo. Elevated chitin inhibited formation of normal hyphae inside hosts but did not inhibit uptake by macrophages. The subsequent survival and proliferation of the *apm4Δ/Δ* mutant within macrophages and in zebrafish suggest a possible mechanism for non-hyphal forming Candida to proliferate but remain avirulent within host organisms

### Experimental Procedures

#### Candida culture and strains

*C. albicans* yeast were grown overnight in YPD medium with 80μg/mL L-uridine. Overnight cultures were refreshed in new media for 30 minutes at 30°C. Cells were washed three times in PBS, counted and used for infections as described for each experiment. For imaging wild-type and *apm4Δ/Δ* Candida strains in mouse macrophages and zebrafish, cytoplasmic Eno1 was tagged with GFP. Tagged strains were analysed to ensure they had the same phenotypes as untagged wt and *apm4Δ/*Δ cells. The following strains were used:

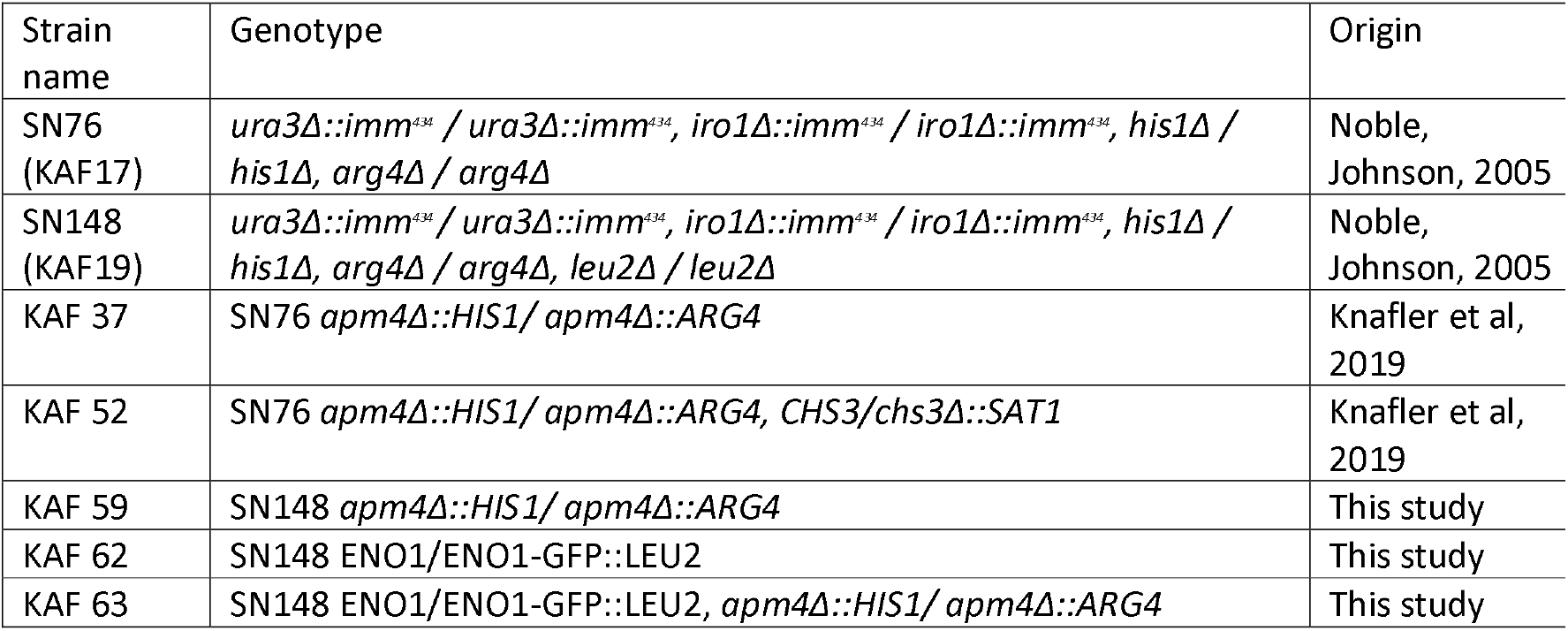
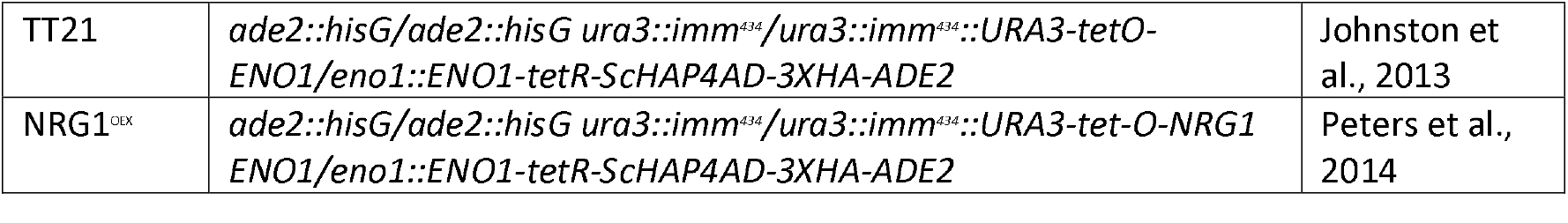

#### Mammalian cell culture and infection

J774 murine macrophage-like cells were maintained in DMEM, with 10% Foetal Bovine Serum, 1% L-glutamine, 1% Penicillin Streptomycin with low glucose for up to 16 passages post thawing. Confluent J774 flasks were used to seed cells at a 5 × 10^5^ cells/mL density in plastic 24 well plates. The day of the experiment the cell health and confluency was checked and the cell media was changed to serum free DMEM.

#### C. albicans immunostaining

Overnight cultures were incubated in fresh YPD medium for 4 hours. After the incubation cells were collected and washed 3 times with PBS. For cell wall chitin staining the cells were resuspended in 1 mL of PBS and 1 μL of a stock 1 mg/mL of Calcofluor white (CFW) (Merck) was added to the suspension. The cells were incubated at room temperature for 5 minutes before being imaged at 150x on a wide field fluorescent microscope. For exposed β glucan staining after the incubation the cultures were centrifuged at 1000 rpm for 3 minutes, cells were resuspended in 1 mL of 4% PFA for 30 minutes and were kept on ice. After the fixation, step cells were washed 3 times with PBS and resuspended in 5 μg/mL of Fc-Dectin from a stock of 2 mg/mL for 1 hr 4°C (Fc-dectin a kind gift from Prof. Gordon Brown, University of Exeter). Cells were washed with freshly made FACs buffer (0.5% bovine serum albumin, 5mM EDTA, 2Mm Na azide in PBS) then resuspended in secondary antibody anti-human IgG conjugated with Alexa-Fluor 488 (Jackson Immuno Research Laboratories) suspended in FACs buffer. The final concentration of the antibody was 1.25 μg/mL. Cells were incubated for 45 minutes on ice in the dark. Following incubation cells were washed 3x with FACs buffer before being imaged (Bain et al., 2014).

#### Phagocytosis of C. albicans

A 1×10^6^ cells/mL suspension of J774 macrophages was seeded on a sterile glass coverslip inside a 24-well plate and incubated overnight. Candida overnight cultures were incubated for 30 minutes in fresh media before being washed and counted to create a Candida cell suspension of 1×10^7^ cells/ml. The macrophage media was changed to serum free media and then they were infected at a 1:10 macrophage to Candida ratio. Cells were incubated 30 minutes in a 37°C incubator with 4% CO2. After the incubation the cells were washed with PBS fixed with 4% PFA (Thermo Scientific) at room temperature for 10 minutes. The slides were washed with PBS and distilled water. The coverslip was attached to the glass slide using 6 μL of mowoil. The slides were imaged with the microscope set at 60x magnification and capturing a Z stack distance of 10 μm slicing at every 1 μm. The images were analysed to determine the percentage of macrophages that contained Candida and the number of *C. albicans* cells per macrophage.

#### Analysis of Candida macrophage Interactions

The microscope chamber was pre-warmed to 37°C with 4% CO2, prepared J774 cultures on 24 well plates were infected with washed cell suspension of *C. albicans* 5 × 10^4^ cells/mL and the timelapse experiment started. Imaging was at Phase 20x magnification using the Widefield Fluorescence microscopy system by Nikon. The microscope captured images every 10 minutes for an overall interaction of 18 hours. The timelapses were used for manual analysis of interaction phenotypes.

#### Exposed dectin-1 levels during macrophage interactions

Macrophages were infected with *C. albicans* for 30-minutes before being fixed and stained. Macrophages were permeabilised with 40 μL per well of 0.25% Triton X 1000 for 3 minutes. After the permeabilization step the cells were washed 3 times with PBS and then the *C. albicans* immunofluorescence staining protocol above was followed. The cells were washed before imaged at 40x using a fluorescence microscope. The data from the images were then collected using the FIJI software by manually drawing around the *Candida* cell periphery to measure using the software the integrated intensity for each channel. The controls for these experiments were cells that were not incubated with macrophages and cells which were incubated with macrophages but not phagocytosed.

#### Zebrafish survival and fungal burden

The zebrafish handling and injection was followed according to the method previously described by (Bojarczuk et al., 2016). 500 Colony forming units of washed overnight cultures of C. albicans was used to inject 1dpf embryos in the caudal vein. The control for the injections was clear PVP + 20% Phenol red. Each injection group was composed of 30 embryos per replicate, the number of biological replicates was 3. Zebrafish survival was monitored up to 4 dpi and the number of deaths was recorded every 24 hours. The death of a zebrafish was determined by cessation of heartbeat. Survival curves were generated in Graphpad Prism for each replicate; statistical test, Mantel-Cox survival test.

The fungal burden studied through fluorescence involved zebrafish infection with C. albicans, the embryos were imaged daily up to 3 dpi. The microscope was capturing Z stacks of the zebrafish embryos at 10x magnification. The maximum intensity projections of the Z stacks and the images of zebrafish were analysed using the FIJI software by drawing around the perimeter of the zebrafish and quantifying the integrated intensity. The results were normalised to uninfected zebrafish from the same experiment. The results were analysed in the Prism software using a one-way ANOVA statistical test.

#### Fixing and imaging fixed zebrafish embryos

Nacre Zebrafish embryos at 1 dpf were injected with 500 cfu of *C. albicans* expressing cytoplasmic GFP, as a control zebrafish were injected with PVP + 20% Phenol red. At 1 dpi about 10 zebrafish embryos per group were fixed in an Eppendorf tube with 4% PFA (Thermo Scientific) overnight at 4°C. The next day PFA was removed, and the embryos were washed twice for 2 minutes with PBS-Tween (Sigma) (1:200 dilution of 20% Tween20). For the imaging step the fixed embryos were mounted on imaging plates and were imaged using the 40x water lens of the confocal. The microscope was set up for imaging the GFP channel.

#### Quantification of single cell observations

The proliferation timelapse experiments were analysed by observations of every macrophage and the Candida cells inside it throughout the course of the timelapse. Observations about how the Candida cells looked before uptake and the changes that occurred during the interaction were recorded. The timepoints of those changes were also recorded. The results were then processed and presented as proportions using Prism software. Only changes inside the phagolysosome were recorded.

The Measure function in FIJI software was used for the quantification of fluorescence to assess different objects (cells) in images. Area, mean intensity and integrated intensity were recorded. The process was manual as shapes were drawn around the periphery of cells to create a closed object.

#### Statistical Analysis

The statistical tests used in this study were a Mann-Whitney test, Fisher’s exact test, One-way ANOVA and a Mantel-Cox survival test, unless otherwise stated. The stars in figures represented the significance values with a confidence interval of 95%. No significance was illustrated by ns and a p value of more than 0.05; * p value of 0.05-0.01; ** indicated significance p value 0.01-0.001; *** indicate a p value of 0.001 <0.0001; **** p value is less or equal to 0.0001. GraphPad Prism version 9 was used for all the results presented.

## Acknowledgements

We thank aquarium staff at the Bateson Centre, Sheffield, UK for their zebrafish husbandry. We thank Prof. Gordon Brown MRC Centre for Medical Mycology, University of Exeter, UK for the kind gift of Fc-dectin. We thank Professor Robert Wheeler, University of Maine, USA for the kind gift of the TT21 and NRG1^oex^ strains.

## Author Contributions

Conceptualization, SAJ, KRA; Formal analysis, SC; Funding acquisition, KA, SAJ; Investigation, SC; Methodology, SAJ, KRA; Project administration, KA, SAJ; Resources, SAJ, KRA; Supervision, SAJ, KRA; Validation, SAJ; Writing – original draft, SC, KRA, SAJ; Writing – review & editing, SC, KRA, SAJ

## Funding

This research was supported by a BBSRC White Rose DTP studentship to SC BB/M011151/1. SAJ was supported by Medical Research Council and Department for International Development Career Development Award Fellowship MR/J009156/1. SAJ was additionally supported by a Krebs Institute Fellowship, and Medical Research Council Centre grant G0700091. Work was performed in the Wolfson Laboratories for Zebrafish Models of Infection funded by the Wolfson Foundation and Royal Society WLR\R1\170024.

## Notes

### Competing Interest Statement

The authors have declared no competing interest.

